# The impact of viral mutations on recognition by SARS-CoV-2 specific T-cells

**DOI:** 10.1101/2021.04.08.438904

**Authors:** Thushan I. de Silva, Guihai Liu, Benjamin B Lindsey, Danning Dong, Dhruv Shah, Alexander J. Mentzer, Adrienn Angyal, Rebecca Brown, Matthew D. Parker, Zixi Ying, Xuan Yao, Lance Turtle, Susanna Dunachie, COVID-19 Genomics UK (COG-UK) Consortium, Mala K. Maini, Graham Ogg, Julian C. Knight, Yanchun Peng, Sarah L. Rowland-Jones, Tao Dong

## Abstract

We identify amino acid variants within dominant SARS-CoV-2 T-cell epitopes by interrogating global sequence data. Several variants within nucleocapsid and ORF3a epitopes have arisen independently in multiple lineages and result in loss of recognition by epitope-specific T-cells assessed by IFN-γ and cytotoxic killing assays. These data demonstrate the potential for T-cell evasion and highlight the need for ongoing surveillance for variants capable of escaping T-cell as well as humoral immunity.

## Main

Evolution of SARS-CoV-2 can lead to evasion from adaptive immunity generated following infection and vaccination. Much focus has been on humoral immunity and spike protein mutations that impair the effectiveness of neutralizing monoclonal antibodies and polyclonal sera. T-cells specific to conserved proteins play a significant protective role in respiratory viral infections such as influenza, particularly in broad heterosubtypic immunity^1^. T-cell responses following SARS-CoV-2 infection are directed against targets across the genome and may play a role in favourable outcomes during acute infection and in immunosuppressed hosts with deficient B-cell immunity^2-4^. While CD8+ T-cells may not provide sterilising immunity, they can protect against severe disease and limit risk of transmission, with a potentially more important role in the setting of antibody escape.

Little is known about the potential for SARS-CoV-2 mutations to impact T-cell recognition. Escape from antigen-specific CD8+ T-cells has been studied extensively in HIV-1 infection, where rapid intra-host evolution renders T-cell responses ineffective within weeks of acute infection^5^. While these escape variants play an important role in the dynamics of chronic viral infections, the opportunities for T-cell escape in acute respiratory viral infections are fewer and consequences are different. Nevertheless, several cytotoxic T-lymphocyte (CTL) escape variants have been described in influenza, such as the R384G substitution in the HLA B*08:01-restricted nucleoprotein_380-388_ and B*27:05-restricted nucleoprotein_383-391_ epitopes^6^. Long-term adaptation of influenza A/H3N2 has been demonstrated, with the loss of one CTL epitope every three years since its emergence in 1968^7^.

To explore the potential for viral evasion from SARS-CoV-2-specific T-cell responses, we conducted a proof-of-concept study, focusing initially on identifying common amino acid mutations within experimentally proven T-cell epitopes and testing the functional implications in selected immunodominant epitopes that we and others have described previously. We conducted a literature review in PubMed and Scopus databases (29^th^ of November 2020; Supplementary Information) that identified 14 publications defining 360 experimentally proven CD4+ and CD8+ T-cell epitopes^2,8-20^. Of these, 53 that were described in ≥1 publication were all CD8+ epitopes (Table S1) and distributed across the genome (n=14 ORF1a, n=5 ORF1b, n=18 S, n=2 M, n=8 N, n=5 ORF3a, n=1 ORF7a). In total 7538 amino acid substitutions or deletions were identified within the 360 T-cell epitopes by searching the COVID-19 Genomics UK consortium (COG-UK) global alignment, dated 29^th^ January 2021 and containing 309,119 sequences (Figure S1, Table S2). 1087 amino acid variants were present within the 53 CD8+ T-cell epitopes with responses described across multiple cohorts, with at least one variant in all epitopes (Figure S2, Table S3).

We focused on evaluating the functional impact of variants within seven immunodominant epitopes (five CD8+, two CD4+) described in our study of UK convalescent donors (Figure 1A)^2^. Of these, all five CD8+ epitopes have been described in at least one other cohort. In particular, responses to the A*03:01/A*11:01-restricted nucleocapsid KTFPPTEPK_361-369_^2,8,10,20^ and A*01:01-restricted ORF3a FTSDYYQLY_207-215_^2,8,10,15^ epitopes are consistently dominant and of high magnitude. We tested the functional avidity of SARS-CoV-2 specific CD4+ and CD8+ polyclonal T-cell lines by interferon (IFN)-γ ELISpots using wild-type and variant peptide titrations (Figure 1B-G). We found that several variants resulted in complete loss of responsiveness to the T-cell lines evaluated: the Q213K variant in the A*01:01-restricted CD8+ ORF3a epitope FTSDYYQLY_207-215_ ^2,8,10,15^, the P13L, P13S and P13T variants in the B*27:05-restricted CD8+ nucleocapsid epitope QRNAPRITF_1-17_ ^2,13^, and T362I and P365S variants in the A*03:01/A*11:01-restricted CD8+ nucleocapsid epitope KTFPPTEPK_361-369_^2,8,10,20^ (Figure 1B-D).

**Figure 1.**
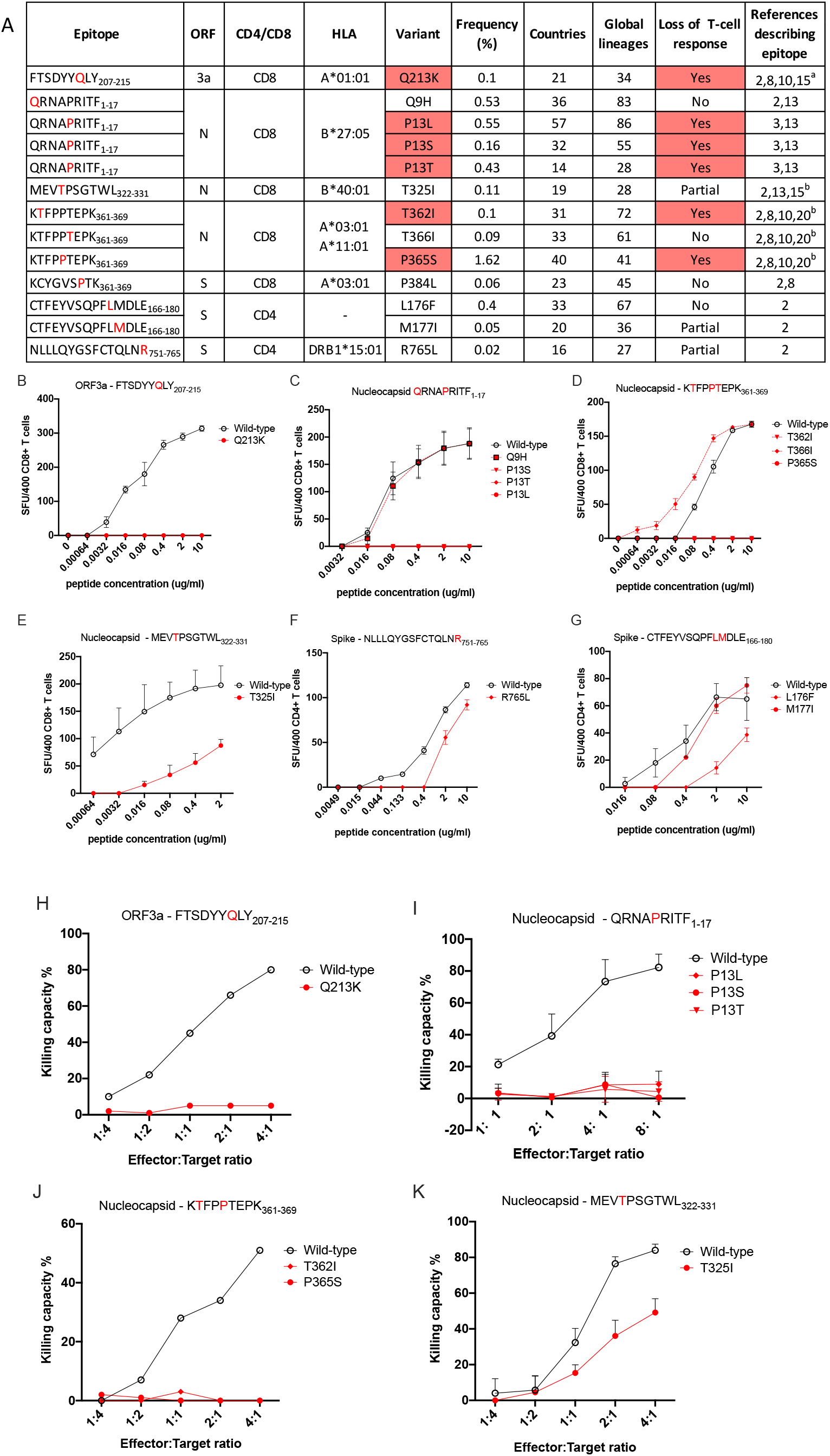
Functional impact of mutations in key SARS-CoV-2 dominant epitopes. **A**. Epitopes and variants studied. Mutated positions detailed in red within wild-type epitope sequence. Frequency indicates % of sequences where variant is seen within COG-UK Global alignment (309,119 sequenced, 29^th^ Jan 2021). Global Lineages refers to Pango lineage assignment. ORF=Open Reading Frame, HLA=Human Leukocyte Antigen. ^a^responses to longer peptide also seen in^18^; ^b^responses to longer peptide also seen in ^10,18^ **B-G**. Recognition of wild-type (black) and mutant (red) peptide titrations by bulk epitope-specific T-cell lines in IFN-γ ELISpot assays. SFU=Spot Forming Units. **H-K**. Ability of CD8+ T-cell lines to kill autologous B-cells loaded with wild-type (black) or mutant (red) peptides in carboxyfluoroscein succinimidyl ester (CFSE) assays. Effector:target ratio denotes proportion of CD8+ T-cell:B-cells in each assay.

In contrast, Q9H in QRNAPRITF_1-17,_ T366I in KTFPPTEPK_361-369,_ P384L in the A*03:01-restricted CD8+ spike epitope KCYGVSPTK_378-386_^2,8^ and M177I in the CD4+ spike epitope CTFEYVSQPFLMDLE_166-180_ ^2^ showed no impact on T-cell recognition (Figures 1C, D, G, S3). Several other variants showed partial loss of T-cell responsiveness, with lower avidity observed to the variant peptide compared to wild-type peptide. These included T325I in the B*40:01-restricted nucleocapsid epitope MEVTPSGTWL_322-331_^2,13,15^, R765L in the DRB1*15:01-restricted CD4+ spike epitope NLLLQYGSFCTQLNR_751-765_^2^, and M177I in the CD4+ spike epitope CTFEYVSQPFLMDLE ^2^ (Figure 1E-G). In order to confirm our findings, we evaluated the impact of CD8+ T-cell epitope variants on CTL killing of peptide-loaded autologous B-cells. Consistent with the ELISpot data, CTL killing ability was significantly impaired by Q213K in ORF3a FTSDYYQLY_207-215_, P13L, P13S and P13T in nucleocapsid QRNAPRITF_1-17_, and T362I and P365S in nucleocapsid KTFPPTEPK_361-369_ (Figure 1H-J). Partial impairment of killing ability was seen with T325I in MEVTPSGTWL_322-331_ (Figure 1K).

In contrast, Q9H in QRNAPRITF_1-17,_ T366I in KTFPPTEPK_361-369,_ P384L in the A*03:01-restricted CD8+ spike epitope KCYGVSPTK_378-386_^2,8^ and M177I in the CD4+ spike epitope CTFEYVSQPFLMDLE_166-180_ ^2^ showed no impact on T-cell recognition (Figures 1C, D, G, S3). Several other variants showed partial loss of T-cell responsiveness, with lower avidity observed to the variant peptide compared to wild-type peptide. These included T325I in the B*40:01-restricted nucleocapsid epitope MEVTPSGTWL_322-331_^2,13,15^, R765L in the DRB1*15:01-restricted CD4+ spike epitope NLLLQYGSFCTQLNR_751-765_^2^, and M177I in the CD4+ spike epitope CTFEYVSQPFLMDLE_166-180_ ^2^ (Figure 1E-G). In order to confirm our findings, we evaluated the impact of CD8+ T-cell epitope variants on CTL killing of peptide-loaded autologous B-cells. Consistent with the ELISpot data, CTL killing ability was significantly impaired by Q213K in ORF3a FTSDYYQLY_207-215_, P13L, P13S and P13T in nucleocapsid QRNAPRITF_1-17_, and T362I and P365S in nucleocapsid KTFPPTEPK_361-369_ (Figure 1H-J). Partial impairment of killing ability was seen with T325I in MEVTPSGTWL_322-331_ (Figure 1K).

T-cell escape can occur via interrupting several mechanisms: antigen processing, binding of MHC to peptide, or T-cell receptor (TCR) recognition of the MHC-peptide complex. While we did not explicitly establish which of these was responsible in each case, it is likely that any partial impairment of T-cell recognition is due to reduced TCR binding to MHC-peptide. Reasons for complete escape are more difficult to predict. As the anchor residues of peptide-MHC binding in A*03:01/A*11:01-restricted KTFPPTEPK_361-369_ are at positions 2 and 9, T362I (position 2) may impair peptide-MHC binding, while P365S (position 5) may affect a T-cell binding residue. The proline changes (P13L, P13S, P13T) in the B*27:05-restricted QRNAPRITF_1-17_ (position 5) again may be at a key T-cell contact residue. The anchor residues for the A*01:01-restricted FTSDYYQLY_207-215_ are predicted to be at position 3 and 9, with auxiliary anchors at positions 2 and 7, which may explain the impact of the Q213K (position 7) variant. In keeping with this, we see no significant impact of these mutations on the predicted binding affinities of epitope to MHC (Table S4). Despite a modest 4-fold decrease in predicted IC_50_ for Q213K compared to wild-type, FTSDYYKLY_207-215_ is still a strong binder to A*01:01.

*Ex vivo* IFN-γ ELISpots in two A*03:01 and two B*27:05 convalescent donors confirmed loss of responses to variant peptides seen with T-cell lines specific to KTFPPTEPK_361-369_ and QRNAPRITF_1-17_ (Figure S4). Thus, our findings using T-cell lines are representative of the circulating T-cell response to these epitopes and of physiological relevance. Interestingly, one A*03:01 donor had low level responses to P365S and T362I, suggesting that subdominant responses via alternative TCR are possible. Our data are also biased by using T-cell lines generated from donors recruited early in the pandemic and therefore likely infected with ‘wild-type’ viruses^2^. While variants that impair antigen processing or MHC-peptide binding result in irreversible loss of T-cell recognition, CTLs with new TCR repertoires can overcome TCR-mediated escape variants, as has been described in HIV-1 infection^21^.

**Figure 1.**
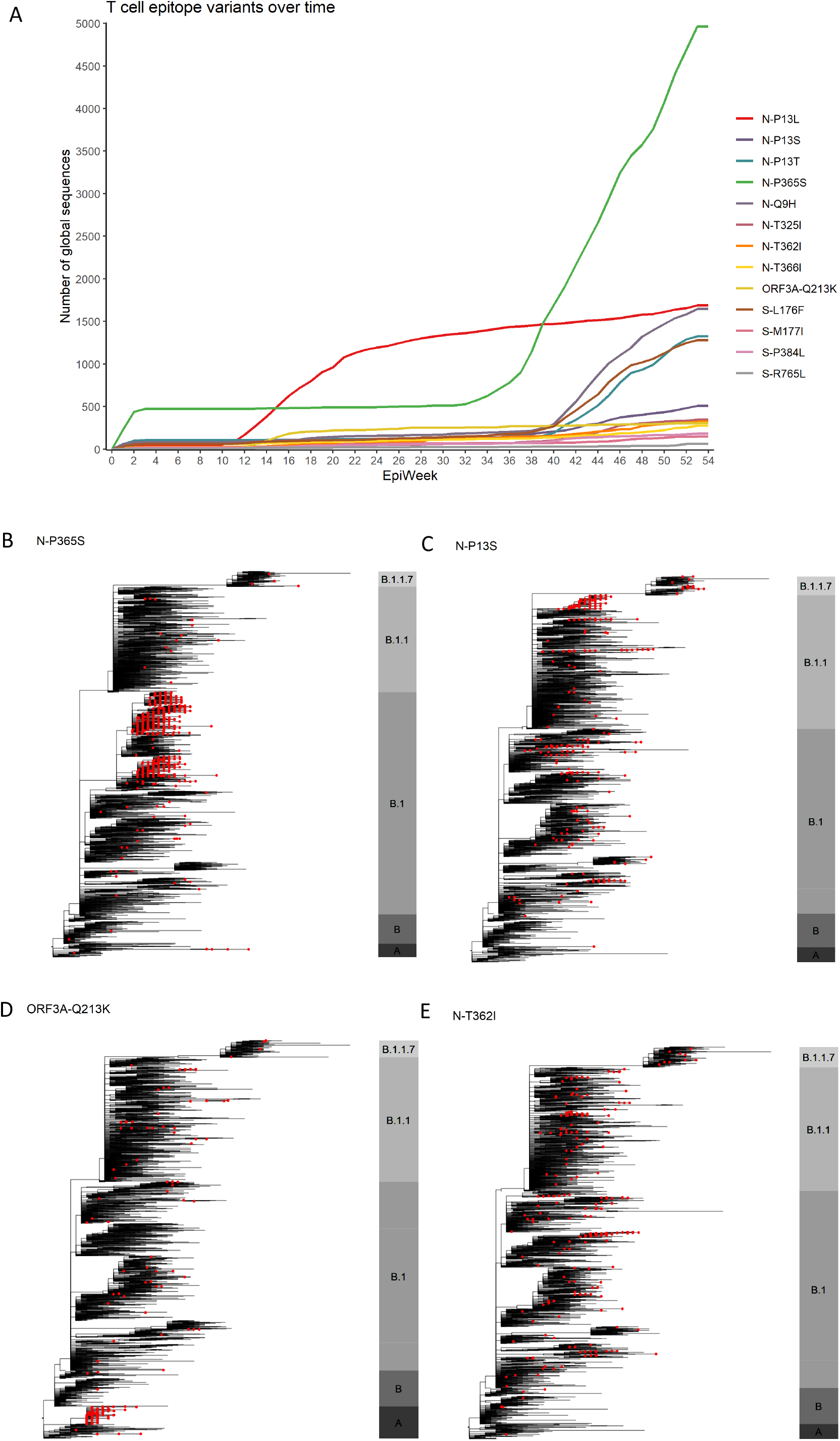
Global presence of variants in key dominant SARS-CoV-2 epitopes. **A**. Weekly frequency over time since beginning of SARS-CoV-2 pandemic of all variants studied in functional experiments. COG-UK global alignment dated 29^th^ Jan 2021 and 309,119 sequences used. Variants named with prefix of SARS-CoV-2 protein (S=spike, N=nucleocapsid), followed by wild-type amino acid, position within protein and variant amino acid. **B-E**. Phylogenies representing global SARS-CoV-2 genomes depicting the presence of epitopes variants impacting T-cell responses. In each case, phylogenies represent all available variant sequences (red tips), along with a selection of non-variant sequences, which were subsampled for visualisation purposes. The bar to the right of each phylogeny is annotated by main ancestral lineages only and not each individual PANGO lineage that viruses belong to. The grapevine pipeline (https://github.com/COG-UK/grapevine) was used for generating the phylogeny based on all data available on GISAID and COG-UK up until 16^th^ of February 2021.

Many variants examined in our study were at relatively low frequency and stable prevalence at the time of writing, other than P365S in KTFPPTEPK_361-369,_ R765L in NLLLQYGSFCTQLNR_751-765_ and variants affecting the proline at position 13 in QRNAPRITF_1-17_ (Figures 1A and 2A). We explored whether variants that result in loss of T-cell recognition appeared as homoplasies in the phylogeny of SARS-CoV-2 suggestive of repeated independent selection, or whether global frequency is due mainly to the expansion of lineages after initial acquisition. While in some cases, variant frequency was dependent on a few successful lineages, P365S, Q213K, T362I, P13L, P13S and P13T had arisen independently on several occasions including within the recently emerged B.1.1.7 lineage (Figures 2B-E, S5A-B). It is important to emphasise that this homoplasy and our functional data do not prove selection due to T-cell escape, which would require demonstration of intra-host evolution. The positions we find important for T-cell recognition may be under selective pressure for reasons other than T-cell immunity. A recent study has documented intra-host evolution of minority variants within A*02:01 and B*40:01 CD8+ epitopes that impair T-cell recognition, though not all epitopes are dominant and very few of the variants studied were represented amongst the global circulating viruses^22^.

There is unlikely to be adequate population immunity at present to see global changes due to T-cell selection akin to what has been seen in adaptation of H3N2 influenza over time^7^. Furthermore, polymorphism in HLA genes restricts the selective advantage of escape within one particular epitope to a relatively small proportion of the population, given the breadth in T-cell responses we and others have shown. Nevertheless, responses to many of the CTL epitopes we have studied are dominant within HLA-matched individuals across many cohorts^2^. As A*03:01, A*11:01 and A*01:01 are common HLA alleles globally, loss of T-cell responses to dominant epitopes such as KTFPPTEPK_361-369_ and FTSDYYQLY_207-215_ may be significant. Substitution of three different amino acid variants at nucleocapsid position 13 within the B*27:05-restricted QRNAPRITF_1-17_ epitope is also striking and suggests significant positive selective pressure at this site. A single dominant, protective B*27:05-restricted epitope has been described in HIV-1 infection, with T-cell escape associated with progression to AIDS. T-cell escape from a B*27:05-restricted influenza A epitope (nucleoprotein_383-391_) has also been observed^6^.

A significant increase in sites under diversifying positive selective pressure was observed around November 2020, most notably in ORF3a, N and S^23^. As vaccine and naturally-acquired population immunity increases further, the frequency of variants we have described should be monitored globally, as well as further changes arising within all immunodominant T-cell epitopes. We have recently incorporated the ability to identify spike T-cell epitope variants in real-time sequence data into the COG-UK mutation explorer dashboard (http://sars2.cvr.gla.ac.uk/cog-uk/). Non-spike T-cell immune responses will also become increasingly important to vaccine-induced immunity as inactivated whole virus vaccines are rolled out. Our findings demonstrate the potential for T-cell evasion and highlight the need for ongoing surveillance for variants capable of escaping T-cell as well as humoral immunity.

## Methods

### Identification of amino acid variants within T-cell epitopes

Variants within the 360 experimentally proven T-cell epitopes were identified using the COVID-19 Genomics UK consortium (COG-UK) global alignment, dated 29^th^ January 2021 and containing 309,119 sequences. Sequences were excluded if they did not contain a start and/stop codon at the beginning and end of each open reading frame (ORF). Each sequence was translated and compared to reference (MN908947.3) using custom python scripts (Python 3.7.6) utilising Biopython (version 1.78).

### Peptide titrations using T-cell lines

Polyclonal CD4+ and CD8+ T-cell lines specific for seven previously described immunodominant epitopes^2^ were generated after MHC class I or II tetramer sorting from cultured short-term cultures of SARS-CoV-2 recovered donor peripheral blood mononuclear cells (PBMCs). Antigen-specific T-cells were confirmed by corresponding tetramer staining. The functional avidity of T-cell lines was assessed by IFN-γ ELISpot assays performed as described previously^24^, by stimulation with wild-type and variant peptides starting at 10μg/mL and serial 1:5 dilutions. Peptides were synthesised by GenScript Biotech (Netherlands) B.V. To quantify antigen-specific responses, spots of the control wells were subtracted from test wells and results expressed as spot forming units (SFU) per 10^6^ PBMCs. If negative control wells had >30 SFU/10^6^ PBMCs or positive control (phytohemagglutinin) were negative, results were considered invalid. Duplicate wells were used for each test and results are from three to seven independent experiments.

### Cytotoxic T-lymphocyte (CTL) killing assays

Autologous B-cells were stained with 0.5μmol/L carboxyfluoroscein succinimidyl ester (CFSE, Thermo Fisher Scientific) before wild-type or variant peptide loading at 1μg/mL for one hour. Peptide-loaded B-cells were co-cultured with CTLs at a range of effector:target (E:T) ratios from 1:4 to 8:1 at 37°C for 6 hours and cells stained with 7-AAD (eBioscience) and CD19-BV42 (eBioscience). Assessment of cell death in each condition was based on the CFSE/7-AAD population present.

### Predictions of binding strength of peptides to MHC

NetMHCpan 4.1 (http://www.cbs.dtu.dk/services/NetMHCpan/) was used to predict the binding strength of wild type and variant epitopes under standard settings (strong binder % rank 0.5, weak binder % rank 2). The predicted affinity (IC_50_ nM) for variant epitopes was compared with wild type.

### Phylogenetic tree generation

Phylogenies were generated using the grapevine pipeline (https://github.com/COG-UK/grapevine) based on all data available on GISAID and COG-UK up until 16^th^ February 2021. In order to visualise all sequences with a specific amino acid variant of interest in a global context, a representative sample of global sequences was obtained in two steps. First, one sequence per country per epi week was selected randomly, followed by random sampling of the remaining sequences to generate a sample of 4000 down-sampled sequences. The global tree was then pruned using code adapted from the tree-manip package (https://github.com/josephhughes/tree-manip). The tips of sequences with amino acid variants impacting T-cell recognition were colour-coded. Visualisations were produced using R/ape, R/ggplot2, R/ggtree, R/treeio, R/phangorn, R/stringr, R/dplyr, R/aplot.

### Ex vivo IFN-γ ELISpots in SARS-CoV-2 recovered donors

Cryopreserved PBMCs were used from SARS-CoV-2 recovered donors recruited into the Sepsis Immunomics study with ethical approval from the South Central -Oxford C Research Ethics Committee in England (Ref 13/SC/0149). These were used for *ex vivo* IFN-γ ELISpots with wild-type and variant peptides. Peptides were added to 200,000 PBMCs at a final concentration of 2μg/mL for 16-18 hours (two replicates per condition). Results were interpreted as detailed above. PBMCs used were from samples taken when patients were between 35 to 53 days from symptom onset.

## Acknowledgements

This work is supported by the UK Medical Research Council (MRC); Chinese Academy of Medical Sciences (CAMS) Innovation Fund for Medical Sciences (CIFMS), China; National Institute of Health Research (NIHR) Oxford Biomedical Research Centre and by UK Research and Innovation (UKRI)/NIHR through the UK Coronavirus Immunology Consortium (UK-CIC). Sequencing of SARS-CoV-2 samples and collation of data was undertaken by the COG-UK CONSORTIUM. COG-UK is supported by funding from the Medical Research Council (MRC) part of UK Research & Innovation (UKRI), the National Institute of Health Research (NIHR) and Genome Research Limited, operating as the Wellcome Sanger Institute. TIdS is supported by a Wellcome Trust Intermediate Clinical Fellowship (110058/Z/15/Z). MDP is funded by the NIHR Sheffield Biomedical Research Centre (BRC – IS-BRC-1215-20017). JCK is a Wellcome Investigator (WT204969/Z/16/Z) and supported by NIHR Oxford Biomedical Research Centre and CIFMS. The views expressed are those of the authors and not necessarily those of the NIHR, or MRC.

## Contributions

TIdS and TD conceptualized the project; TD, TIdS and YP designed and supervised T cell experiments, BBL and MDP conducted the viral sequence analyses, DS conducted the literature review and collated T-cell epitope information, GL, DD performed experiments and analysed the data, XY, ZY, AA. and RB provided critical reagents and technical assistance, JCK and AJM, established clinical cohorts; TIdS and TD wrote and edited the original draft, all co-authors reviewed and edited the manuscript.

