## Supplementary information for "The impact of viral mutations on recognition by SARS-CoV-2 specific T-cells"

Search terms used for literature review of SARS-CoV-2 T-cell epitopes in PubMed and Scopus databases

(TITLE-ABS-KEY( ("nCov" OR "Novel Coronavirus" OR "2019 Novel Coronavirus" OR "Covid-19" OR "2019-nCoV" OR "Severe Acute Respiratory Syndrome-Coronavirus-2" OR "SARS-CoV-2")) AND TITLE-ABS-KEY(( "T-cell*" OR "Tcell*" OR "T Cell*" OR "Peptide" OR "cd4+" OR "cd8+" OR "CD8+" OR "CD4+" OR "t-lymphocyt*" OR "T Lymphocyte*")) AND TITLE-ABS-KEY("Epitope*"))

Reasons for rejection of 271/285 publications identified

1. Did not investigate SARS-CoV-2 T-cell responses on convalescent donors (included B-cell and anitbody responses, responses to other viruses e.g. SARS, MERS). N = 53.
2. SARS-CoV-2 T-cell responses explored with whole protein and/or overlapping peptides, with no individual epitope-specific data. N = 31.
3. SARS-CoV-2 T-cell epitopes described soley with bioinformatic prediction (not experimentally proven). N = 129.
4. Reviews of SARS-CoV-2 T-cell responses. N = 44.
5. SARS-CoV-2 T-cell responses in non-human studies. N = 6.
6. Other. N = 7.

Of note, while some publications had defined optimal T-cell epitopes and HLA restriction, some simply reported the sequence of the longer overlapping peptides containing potential epitope to which responses were seen.

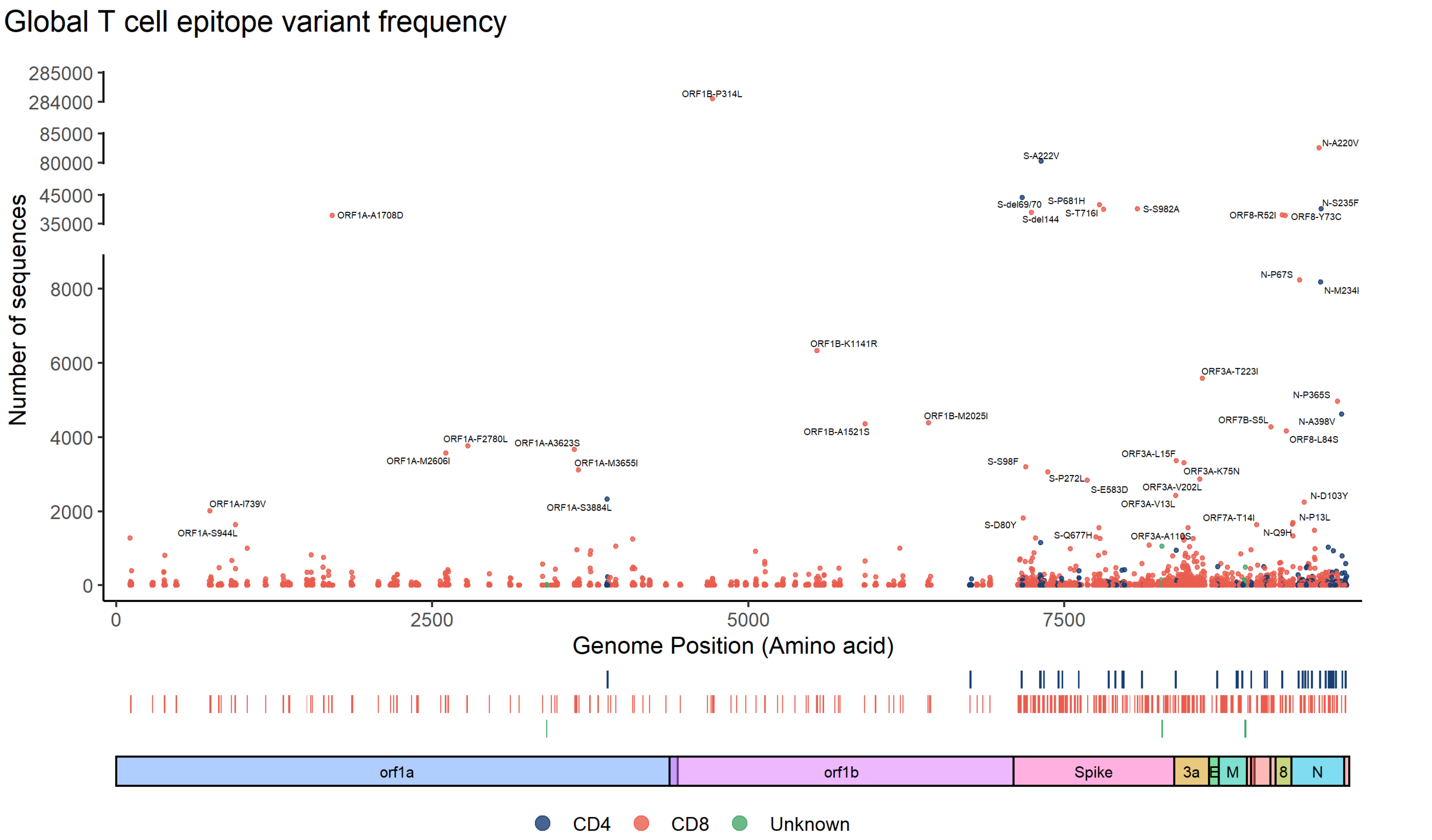


**Figure S1. Frequency of amino acid variants within experimentally proven T-cell epitopes identified in the literature.** Variants within 360 epitopes identified by searching the COVID-19 Genomics UK Consortium (COG-UK) global alignment dated 19th January 2021 (309,119 sequences). CD4 and CD8 epitopes are coloured in blue and red respectively. Open reading frame (ORF) and variant annotated for mutations of higher frequency following convention of wild type amino acid, followed by amino acid position within each ORF and replacement amino acid (e.g. P681H for a proline to histidine at position 681).

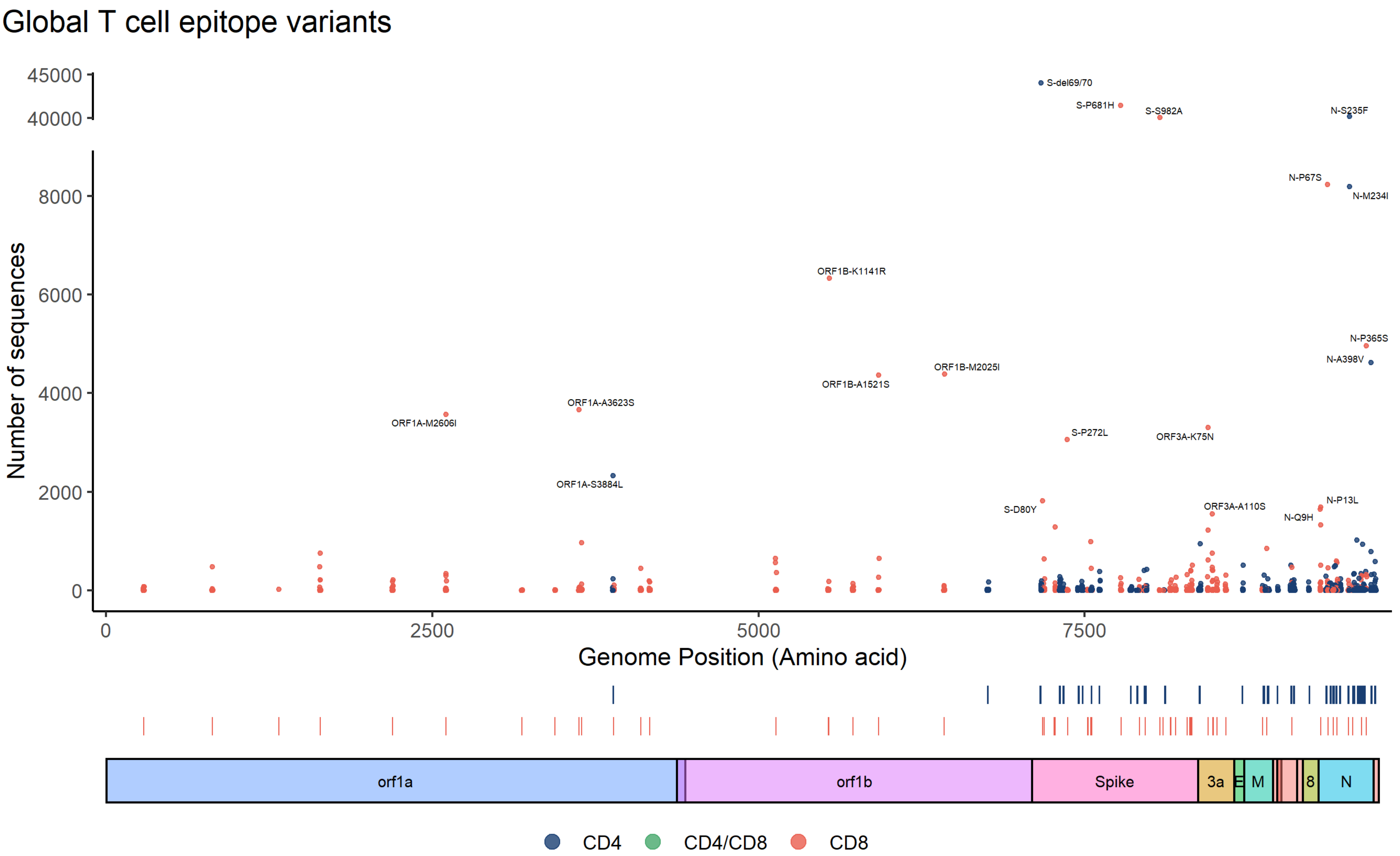


**Figure S2. Frequency of amino acid variants within a focused set of experimentally proven T-cell epitopes identified in the literature.** Variants in epitopes identified by searching the COVID-19 Genomics UK Consortium (COG-UK) global alignment dated 19th January 2021 (309,119 sequences). Shown are variants within CD8+ T-cells epitopes described in two or more cohorts (n=53), as well as all experimentally proven CD4+ T-cell epitopes (N=53). CD4 and CD8 epitopes are coloured in blue and red respectively. Open reading frame (ORF) and variant annotated for mutations of higher frequency following convention of wild type amino acid, followed by amino acid position within each ORF and replacement amino acid (e.g. P681H for a proline to histidine at position 681).

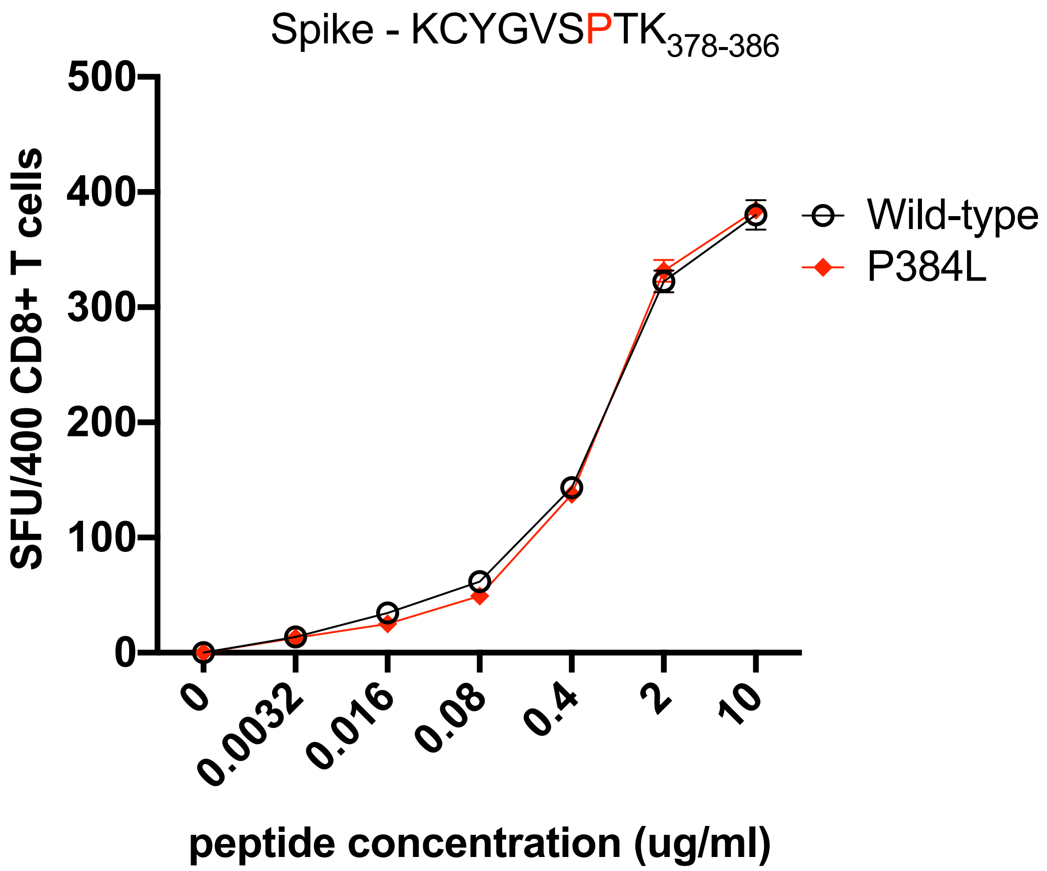


**Figure S3.** IFN-γ ELISpot responses to CD8+ T-cell line specific to the A*03:01-restricted spike epitope $KCYGVSPTK_{378\text{-}386}$ using wild-type and P384L mutant peptide titrations (n=3 replicates per condition). SFU = spot forming units. Mean and standard deviation displayed.

| Epitope | ORF | Amin acid residues | HLA restriction | Variant | Loss of T-cell response | Predicted binding affinity to MHC (IC50 nM) | Difference in predicted binding from wild-type epitope | Binding level |
|---|---|---|---|---|---|---|---|---|
| FTSDYYQLY | 3a | 207 - 215 | A*01:01 | - | - | 5.36 | - | Strong |
| FTSDYYKLY | | | | Q213K | Yes | 21.36 | 3.99-fold decrease | Strong |
| QRNAPRITF | N | 1 - 17 | B*27:05 | - | - | 509.43 | - | Strong |
| QRNALRITF | | | | P13L | Yes | 83.06 | 6.1-fold increase | Strong |
| QRNASRITF | | | | P13S | Yes | 346.15 | 1.47-fold increase | Strong |
| QRNATRITF | | | | P13T | Yes | 229.47 | 2.22-fold increase | Strong |
| MEVTPSGTWL | N | 322 - 331 | B*40:01 | - | - | 33.85 | - | Strong |
| MEVIPSGTWL | | | | T325I | Partial | 34.59 | Neutral | Strong |
| KTFPPTEPK | N | 361 - 369 | A*03:01 | - | - | 19.39 | - | Strong |
| KIFPPTEPK | | | | T362I | Yes | 16.20 | Neutral | Strong |
| KTFPSTEPK | | | | P365S | Yes | 24.96 | Neutral | Strong |
| KTFPPTEPK | | | A*11:01 | - | - | 8.51 | - | Strong |
| KIFPPTEPK | | | | T362I | Yes | 11.12 | Neutral | Strong |
| KTFPSTEPK | | | | P365S | Yes | 7.73 | Neutral | Strong |

**Table S4. Binding predictions of wild-type and mutant epitopes to Major Histocompatibility Complex (MHC) alleles.** Predictions derived from NetMHCpan 4.1 (http://www.cbs.dtu.dk/services/NetMHCpan/). ORF = open reading frame, HLA = human leukocyte antigen.

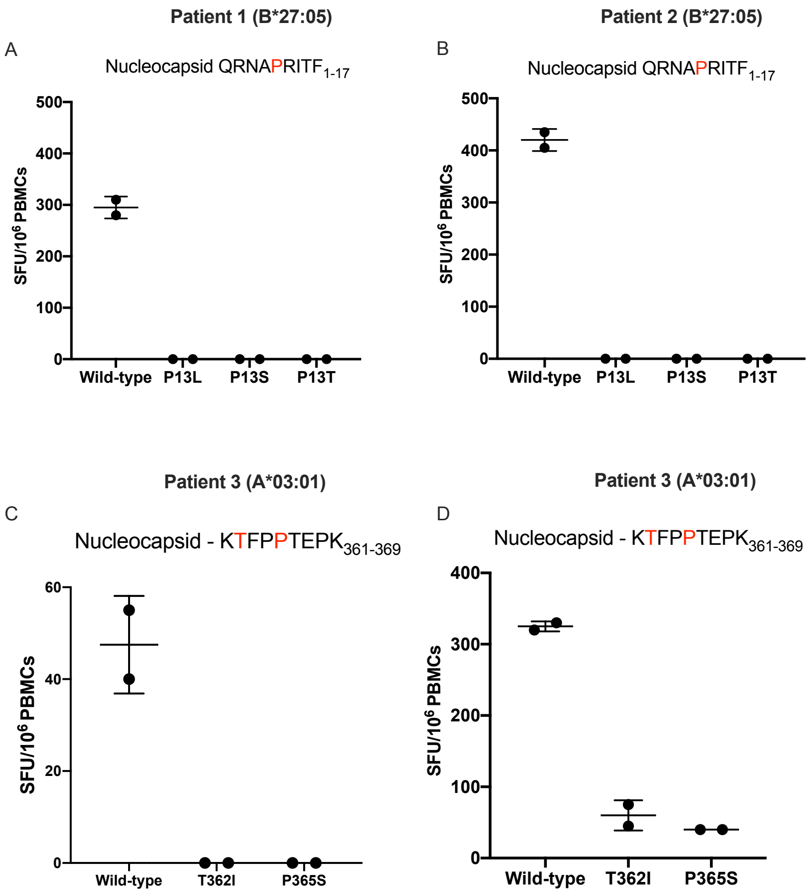


**Figure S4. Ex-vivo IFN-γ ELISpot responses to wild-type and variant peptides.** A and B. Peripheral blood mononuclear cells from two B*27:05 patients tested with the wild-type peptide $QRNAPRITF_{1-17}$ and mutants P13L, P13S and P13T. C and D. Peripheral blood mononuclear cells from two A*03:01 patients tested with wild-type peptide $KTFPPTEPK_{361-369}$ and mutant T362I and P365S.

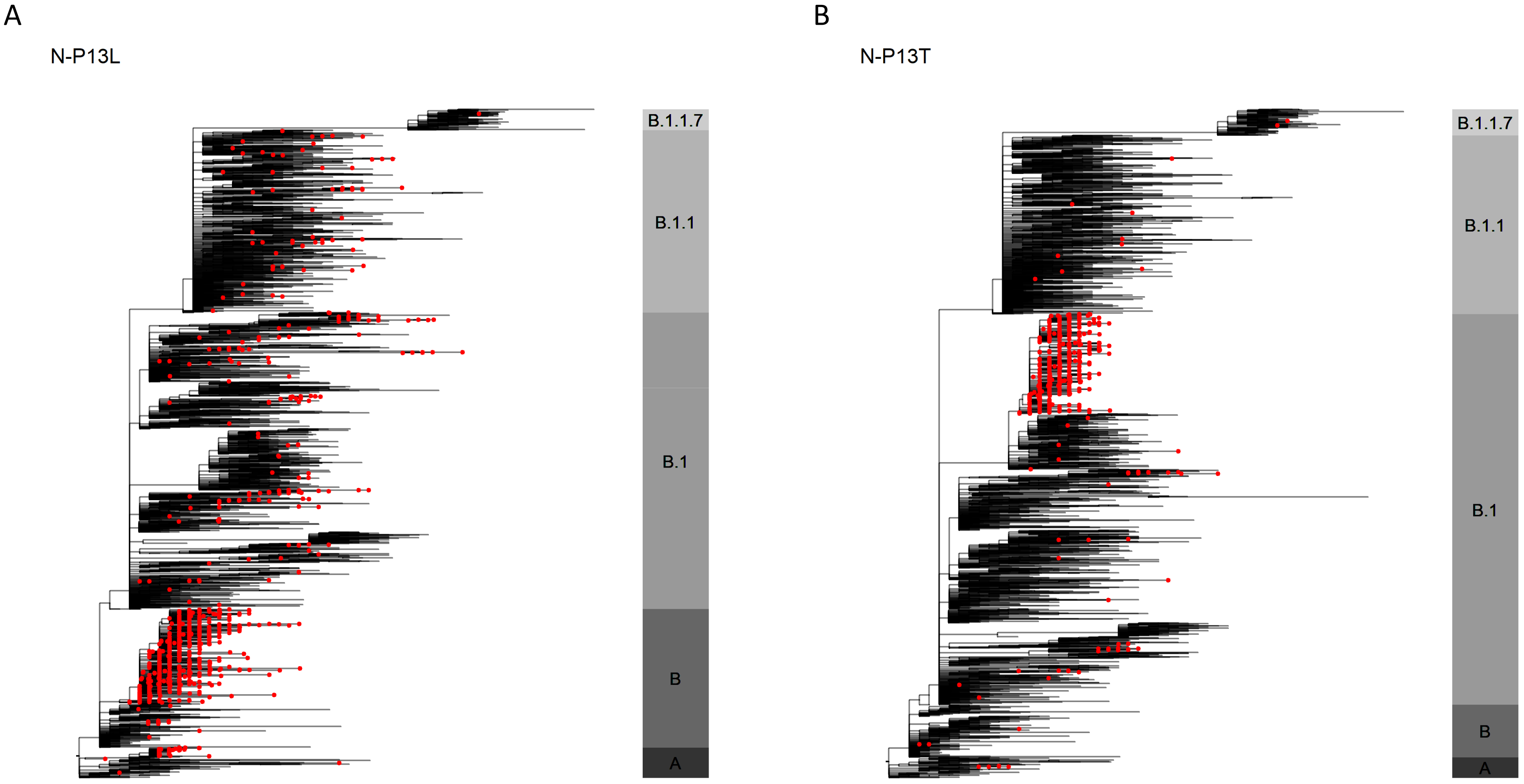


**Figure S5. A representative phylogenetic trees of global SARS-CoV-2 genomes depicting the presence of P13L (A) and P13T (B) variants in the nucleocapsid QRNAPRITF$_{1\text{-}17}$ CD8+ T-cell epitope.** Phylogenies represent all available P13L/P13T sequences (red tips), along with a selection of non-P13L/P13T sequences, which were subsampled for visualisation purposes. The bar to the right of each phylogeny is annotated by main ancestral lineages only and not each individual PANGO lineage that viruses belong to. The grapevine pipeline (https://github.com/COG-UK/grapevine) was used for generating the phylogeny based on all data available on GISAID and COG-UK up until 16$^{th}$ of February 2021.

### Supplementary Authors

**Funding acquisition, Leadership and supervision, Metadata curation, Project administration, Samples and logistics, Sequencing and analysis, Software and analysis tools, and Visualisation:**
Dr Samuel C Robson PhD [13].

**Funding acquisition, Leadership and supervision, Metadata curation, Project administration, Samples and logistics, Sequencing and analysis, and Software and analysis tools:**
Prof Nicholas J Loman PhD [41] and Dr Thomas R Connor PhD [10, 69].

**Leadership and supervision, Metadata curation, Project administration, Samples and logistics, Sequencing and analysis, Software and analysis tools, and Visualisation:**
Dr Tanya Golubchik PhD [5].

**Funding acquisition, Metadata curation, Samples and logistics, Sequencing and analysis, Software and analysis tools, and Visualisation:**
Dr Rocio T Martinez Nunez PhD [42].

**Funding acquisition, Leadership and supervision, Metadata curation, Project administration, and Samples and logistics:**
Dr Catherine Ludden PhD [88].

**Funding acquisition, Leadership and supervision, Metadata curation, Samples and logistics, and Sequencing and analysis:**
Dr Sally Corden PhD [69].

**Funding acquisition, Leadership and supervision, Project administration, Samples and logistics, and Sequencing and analysis:**
Ian Johnston [99] and Dr David Bonsall PhD [5].

**Funding acquisition, Leadership and supervision, Sequencing and analysis, Software and analysis tools, and Visualisation:**
Prof Colin P Smith PhD [87] and Dr Ali R Awan PhD [28].

**Funding acquisition, Samples and logistics, Sequencing and analysis, Software and analysis tools, and Visualisation:**
Dr Giselda Bucca PhD [87].

**Leadership and supervision, Metadata curation, Project administration, Samples and logistics, and Sequencing and analysis:**
Dr M. Estee Torok FRCP [22, 101].

**Leadership and supervision, Metadata curation, Project administration, Samples and logistics, and Visualisation:**
Dr Kordo Saeed MD/ FRCPath [81, 110] and Dr Jacqui A Prieto PhD [83, 109].

**Leadership and supervision, Metadata curation, Project administration, Sequencing and analysis, and Software and analysis tools:**
Dr David K Jackson PhD [99].

**Metadata curation, Project administration, Samples and logistics, Sequencing and analysis, and Software and analysis tools:**
Dr William L Hamilton PhD [22].
**Metadata curation, Project administration, Samples and logistics, Sequencing and analysis, and Visualisation:**
Dr Luke B Snell MSc/ MBBS [11].

**Funding acquisition, Leadership and supervision, Metadata curation, and Samples and logistics:**
Dr Catherine Moore [69].

**Funding acquisition, Leadership and supervision, Project administration, and Samples and logistics:**
Dr Ewan M Harrison PhD [99, 88].

**Leadership and supervision, Metadata curation, Project administration, and Samples and logistics:**
Dr Sonia Goncalves PhD [99].

**Leadership and supervision, Metadata curation, Samples and logistics, and Sequencing and analysis:**
Prof Ian G Goodfellow PhD [24], Dr Derek J Fairley PhD [3, 72], Prof Matthew W Loose PhD [18] and Joanne Watkins MSc [69].

**Leadership and supervision, Metadata curation, Samples and logistics, and Software and analysis tools:**
Rich Livett MSc [99].

**Leadership and supervision, Metadata curation, Samples and logistics, and Visualisation:**
Dr Samuel Moses MD [25, 106].

**Leadership and supervision, Metadata curation, Sequencing and analysis, and Software and analysis tools:**
Dr Roberto Amato PhD [99], Dr Sam Nicholls PhD [41] and Dr Matthew Bull PhD [69].

**Leadership and supervision, Project administration, Samples and logistics, and Sequencing and analysis:**
Prof Darren L Smith PhD [37, 58, 105].

**Leadership and supervision, Sequencing and analysis, Software and analysis tools, and Visualisation:**
Dr Jeff Barrett PhD [99] and Prof David M Aanensen PhD [14, 114].

**Metadata curation, Project administration, Samples and logistics, and Sequencing and analysis:**
Dr Martin D Curran PhD [65], Dr Surendra Parmar PhD [65], Dr Dinesh Aggarwal MRCP [95, 99, 64] and Dr James G Shepherd MBChB/MRCP [48].

**Metadata curation, Project administration, Sequencing and analysis, and Software and analysis tools:**
Dr Matthew D Parker PhD [93].

**Metadata curation, Samples and logistics, Sequencing and analysis, and Visualisation:**
Dr Sharon Glaysher PhD [61].

**Metadata curation, Sequencing and analysis, Software and analysis tools, and Visualisation:**
Dr Matthew Bashton PhD [37, 58], Dr Anthony P Underwood PhD [14, 114], Dr Nicole Pacchiarini PhD [69] and Dr Katie F Loveson PhD [77].

**Project administration, Sequencing and analysis, Software and analysis tools, and Visualisation:**
Dr Alessandro M Carabelli PhD [88].

**Funding acquisition, Leadership and supervision, and Metadata curation:**
Dr Kate E Templeton PhD [53, 90].

**Funding acquisition, Leadership and supervision, and Project administration:**
Dr Cordelia F Langford PhD [99], John Sillitoe BEng [99], Dr Thushan I de Silva PhD [93] and Dr Dennis Wang PhD [93].

**Funding acquisition, Leadership and supervision, and Sequencing and analysis:**
Prof Dominic Kwiatkowski [99, 107], Prof Andrew Rambaut DPhil [90], Dr Justin O'Grady PhD [70, 89] and Dr Simon Cottrell PhD [69].

**Leadership and supervision, Metadata curation, and Sequencing and analysis:**
Prof Matthew T.G. Holden PhD [68] and Prof Emma C Thomson PhD/FRCP [48].

**Leadership and supervision, Project administration, and Samples and logistics:**
Dr Husam Osman PhD [64, 36], Dr Monique Andersson PhD [59], Prof Anoop J Chauhan [61] and Dr Mohammed O Hassan-Ibrahim PhD/FRCPath [6].

**Leadership and supervision, Project administration, and Sequencing and analysis:**
Dr Mara Lawniczak [99].

**Leadership and supervision, Samples and logistics, and Sequencing and analysis:**
Prof Ravi Kumar Gupta PhD [88, 113], Dr Alex Alderton PhD [99], Dr Meera Chand [66], Dr Chrystala Constantinidou PhD [94], Dr Meera Unnikrishnan PhD [94], Prof Alistair C Darby PhD [92], Prof Julian A Hiscox PhD [92] and Prof Steve Paterson PhD [92].

**Leadership and supervision, Sequencing and analysis, and Software and analysis tools:**
Dr Inigo Martincorena [99], Prof David L Robertson PhD [48], Dr Erik M Volz PhD [39], Dr Andrew J Page PhD [7] and Prof Oliver G Pybus DPhil [23].

**Leadership and supervision, Sequencing and analysis, and Visualisation:**
Dr Andrew R Bassett PhD [99].

**Metadata curation, Project administration, and Samples and logistics:**

Dr Cristina V Ariani PhD [99], Dr Michael H Spencer Chapman MBBS [99, 88], Dr Kathy K Li MBBCh/FRCPath [48], Dr Rajiv N Shah BMBS/MRCP/MSc [48], Dr Natasha G Jesudason MBChB MRCP FRCPath [48] and Dr Yusri Taha MD/PhD [50].

**Metadata curation, Project administration, and Sequencing and analysis:**
Martin P McHugh MSc [53], Dr Rebecca Dewar PhD [53].

**Metadata curation, Samples and logistics, and Sequencing and analysis:**
Dr Aminu S Jahun PhD [24], Dr Claire McMurray PhD [41], Ms Sarojini Pandey MSc [84], Dr James P McKenna PhD [3], Dr Andrew Nelson PhD [58, 105], Dr Gregory R Young PhD [37, 58], Dr Clare M McCann PhD [58, 105] and Mr Scott Elliott [61].

**Metadata curation, Samples and logistics, and Visualisation:**
Ms Hannah Lowe MSc [25].

**Metadata curation, Sequencing and analysis, and Software and analysis tools:**
Dr Ben Temperton Ph.D. [91], Dr Sunando Roy PhD [82], Dr Anna Price PhD [10], Dr Sara Rey PhD [69] and Mr Matthew Wyles [93].

**Metadata curation, Sequencing and analysis, and Visualisation:**
Stefan Rooke MSc [90] and Dr Sharif Shaaban PhD [68].

**Project administration, Samples and logistics, Sequencing and analysis:**
Dr Mariateresa de Cesare PhD [98].

**Project administration, Samples and logistics, and Software and analysis tools:**
Laura Letchford BSc [99].

**Project administration, Samples and logistics, and Visualisation:**
Miss Siona Silveira MSc [81], Dr Emanuela Pelosi FRCPath [81] and Dr Eleri Wilson-Davies MD/FRCPath [81].

**Samples and logistics, Sequencing and analysis, and Software and analysis tools:**
Dr Myra Hosmillo PhD [24].

**Sequencing and analysis, Software and analysis tools, and Visualisation:**
Áine O'Toole MSc [90], Dr Andrew R Hesketh PhD [87], Mr Richard Stark MSc [94], Dr Louis du Plessis PhD [23], Dr Chris Ruis PhD [88], Dr Helen Adams PhD [4] and Dr Yann Bourgeois PhD [76].

**Funding acquisition, and Leadership and supervision:**
Dr Stephen L Michell PhD [91], Prof Dimitris Grammatopoulos PhD/FRCPath [84, 112], Dr Jonathan Edgeworth PhD/FRCPath [12], Prof Judith Breuer MD [30, 82], Prof John A Todd PhD [98] and Dr Christophe Fraser PhD [5].

**Funding acquisition, and Project administration:**
Dr David Buck PhD [98] and Michaela John BSc [9].

**Leadership and supervision, and Metadata curation:**
Dr Gemma L Kay PhD [70].

**Leadership and supervision, and Project administration:**
Steve Palmer [99], Prof Sharon J Peacock [88, 64] and David Heyburn [69].

**Leadership and supervision, and Samples and logistics:**
Danni Weldon BSc [99], Dr Esther Robinson PhD [64, 36], Prof Alan McNally PhD [41, 86], Dr Peter Muir PhD [64], Dr Ian B Vipond PhD [64], Dr John BoYes MBChB [29], Dr Venkat Sivaprakasam PhD [46], Dr Tranprit Saluja FRCPath/MD [75], Dr Samir Dervisevic FRCPath [54] and Dr Emma J Meader FRCPath [54].

**Leadership and supervision, and Sequencing and analysis:**
Dr Naomi R Park PhD [99], Karen Oliver BSc [99], Dr Aaron R Jeffries Ph.D. [91], Dr Sascha Ott PhD [94], Dr Ana da Silva Filipe PhD [48], Dr David A Simpson PhD [72] and Dr Chris Williams MB BS [69].

**Leadership and supervision, and Visualisation:**
Dr Jane AH Masoli MBChB [73, 91].

**Metadata curation, and Samples and logistics:**
Dr Bridget A Knight PhD. [73, 91], Dr Christopher R Jones Ph.D. [73, 91], Mr Cherian Koshy MSc CSci FIBMS [1], Miss Amy Ash BSc [1], Dr Anna Casey PhD [71], Dr Andrew Bosworth PhD [64, 36], Dr Liz Ratcliffe PhD [71], Dr Li Xu-McCrae PhD [36], Miss Hannah M Pymont MSc [64], Ms Stephanie Hutchings [64], Dr Lisa Berry PhD [84], Ms Katie Jones MSc [84], Dr Fenella Halstead PhD [46], Mr Thomas Davis MSc [21], Dr Christopher Holmes PhD [16], Prof Miren Iturriza-Gomara PhD [92], Dr Anita O Lucaci PhD [92], Dr Paul Anthony Randell MBBCh [38, 104], Dr Alison Cox PhD [38, 104], Pinglawathee Madona [38, 104], Dr Kathryn Ann Harris PhD [30], Dr Julianne Rose Brown PhD [30], Dr Tabitha W Mahungu FRCPath [74], Dr Dianne Irish-Tavares FRCPath [74], Dr Tanzina Haque FRCPath PhD [74], Dr Jennifer Hart MRCP [74], Mr Eric Witele MSc [74], Mrs Melisa Louise Fenton DipHE [75], Mr Steven Liggett [79], Dr Clive Graham MD [56], Ms Emma Swindells Bsc [57], Ms Jennifer Collins BSc [50], Mr Gary Eltringham BSc [50], Ms Sharon Campbell MSc [17], Dr Patrick C McClure PhD [97], Dr Gemma Clark PhD [15], Dr Tim J Sloan PhD [60], Mr Carl Jones [15] and Dr Jessica Lynch PhD MBChB [2, 111].

**Metadata curation, and Sequencing and analysis:**
Dr Ben Warne MRCP [8], Steven Leonard PhD [99], Jillian Durham BSc [99], Dr Thomas Williams MD [90], Dr Sam T Haldenby PhD [92], Dr Nathaniel Storey PhD [30], Dr Nabil-Fareed Alikhan PhD [70], Dr Nadine Holmes PhD [18], Dr Christopher Moore PhD [18], Mr Matthew Carlile BSc [18], Malorie Perry MSc [69], Dr Noel Craine DPhil [69], Prof Ronan A Lyons MD [80], Miss Angela H Beckett MSc [13], Salman Goudarzi PhD [77], Christopher Fearn MRes [77], Kate Cook [77], Hannah Dent BSc [77] and Hannah Paul MRes [77].

**Metadata curation, and Software and analysis tools:**
Robert Davies [99].

**Project administration, and Samples and logistics:**

Beth Blane BSc [88], Sophia T Girgis MSc [88], Dr Mathew A Beale PhD [99], Katherine L Bellis [99, 88], Matthew J Dorman [99], Eleanor Drury [99], Leanne Kane [99], Sally Kay [99], Dr Samantha McGuigan [99], Dr Rachel Nelson PhD [99], Liam Prestwood [99], Dr Shavanthi Rajatileka PhD [99], Dr Rahul Batra MD [12], Dr Rachel J Williams PhD [82], Dr Mark Kristiansen PhD [82], Dr Angie Green PhD [98], Miss Anita Justice MSc [59], Dr Adhyana I.K Mahanama MD [81, 102] and Dr Buddhini Samaraweera MD [81, 102].

**Project administration, and Sequencing and analysis:**
Dr Nazreen F Hadjirin PhD [88] and Dr Joshua Quick PhD [41].

**Project administration, and Software and analysis tools:**
Mr Radoslaw Poplawski BSc [41].

**Samples and logistics, and Sequencing and analysis:**
Leanne M Kermack MSc [88], Nicola Reynolds PhD [7], Grant Hall BS [24], Yasmin Chaudhry BSc [24], Malte L Pinckert MPhil [24], Dr Iliana Georgana PhD [24], Dr Robin J Moll PhD [99], Dr Alicia Thornton [66], Dr Richard Myers [66], Dr Joanne Stockton PhD [41], Miss Charlotte A Williams BSc [82], Dr Wen C Yew PhD [58], Alexander J Trotter MRes [70], Miss Amy Trebes MSc [98], Mr George MacIntyre-Cockett BSc [98], Alec Birchley MSc [69], Alexander Adams BSc [69], Amy Plimmer [69], Bree Gatica-Wilcox MPhil [69], Dr Caoimhe McKerr PhD [69], Ember Hilvers MA [69], Hannah Jones [69], Dr Hibo Asad PhD [69], Jason Coombes BSc [69], Johnathan M Evans MSc [69], Laia Fina [69], Lauren Gilbert A-Levels [69], Lee Graham BSc [69], Michelle Cronin [69], Sara Kumziene-SummerhaYes MSc [69], Sarah Taylor [69], Sophie Jones MSc [69], Miss Danielle C Groves BA [93], Mrs Peijun Zhang MSc [93], Miss Marta Gallis MSc [93] and Miss Stavroula F Louka MSc [93].

**Samples and logistics, and Software and analysis tools:**
Dr Igor Starinskij Msc MRCP [48].

**Sequencing and analysis, and Software and analysis tools:**
Dr Chris J Illingworth PhD [47], Dr Chris Jackson PhD [47], Ms Marina Gourtovaia MSc [99], Gerry Tonkin-Hill [99], Kevin Lewis [99], Dr Jaime M Tovar-Corona PhD [99], Dr Keith James PhD [99], Dr Laura Baxter PhD [94], Dr Mohammad T. Alam PhD [94], Dr Richard J Orton PhD [48], Dr Joseph Hughes PhD [48], Dr Sreenu Vattipally PhD [48], Dr Manon Ragonnet-Cronin PhD [39], Dr Fabricia F. Nascimento PhD [39], Mr David Jorgensen MSc [39], Ms Olivia Boyd MSc [39], Ms Lily Geidelberg MSc [39], Dr Alex E Zarebski PhD [23], Dr Jayna Raghwani PhD [23], Dr Moritz UG Kraemer DPhil [23], Joel Southgate MSc [10, 69], Dr Benjamin B Lindsey MRCP [93] and Mr Timothy M Freeman MPhil [93].

**Software and analysis tools, and Visualisation:**
Jon-Paul Keatley [99], Dr Joshua B Singer PhD [48], Leonardo de Oliveira Martins PhD [70], Dr Corin A Yeats PhD [14], Dr Khalil Abudahab PhD [14, 114], Mr Ben EW Taylor MEng [14, 114] and Mirko Menegazzo [14].

**Leadership and supervision:**
Prof John Danesh [99], Wendy Hogsden MSc [46], Dr Sahar Eldirdiri MBBS MSc FRCPath [21], Mrs Anita Kenyon MSc [21], Dr Jenifer Mason MBBS [43], Mr Trevor I Robinson MSc [43], Prof Alison Holmes MD [38, 103], Dr James Price PhD [38, 103], Prof John A Hartley PhD [82], Dr Tanya Curran PhD [3], Dr Alison E Mather PhD [70], Dr Giri Shankar [69], Dr Rachel Jones [69], Dr Robin Howe [69] and Dr Sian Morgan FRCPath [9].

**Metadata curation:**
Dr Elizabeth Wastenge MD [53], Dr Michael R Chapman PhD [34, 88, 99], Mr Siddharth Mookerjee MPH [38, 103], Dr Rachael Stanley PhD [54], Mrs Wendy Smith [15], Prof Timothy Peto PhD [59], Dr David Eyre PhD [59], Dr Derrick Crook [59], Dr Gabrielle Vernet MBBS [33], Dr Christine Kitchen PhD [10], Huw Gulliver [10], Dr Ian Merrick PhD [10], Prof Martyn Guest PhD [10], Robert Munn BSc [10], Dr Declan T Bradley [63, 72] and Dr Tim Wyatt [63].

**Project administration:**
Dr Charlotte Beaver [99], Luke Foulser [99], Sophie Palmer [88], Carol M Churcher [88], Ellena Brooks MA [88], Kim S Smith [88], Dr Katerina Galai PhD [88], Georgina M McManus BSc [88], Dr Frances Bolt PhD [38, 103], Dr Francesc Coll PhD [19], Lizzie Meadows MA [70], Dr Stephen W Attwood PhD [23], Dr Alisha Davies [69], Elen De Lacy MSc [69], Fatima Downing [69], Sue Edwards [69], Dr Garry P Scarlett PhD [76], Mrs Sarah Jeremiah MSc [83] and Dr Nikki Smith PhD [93].

**Samples and logistics:**
Danielle Leek Bsc [88], Sushmita Sridhar BS [88, 99], Sally Forrest BSc [88], Claire Cormie [88], Harmeet K Gill PhD [88], Joana Dias MSc [88], Ellen E Higginson PhD [88], Mailis Maes MPhil [88], Jamie Young BSc [88], Michelle Wantoch PhD [7], Sanger Covid Team (www.sanger.ac.uk/covid-team) [99], Dorota Jamrozy [99], Stephanie Lo [99], Dr Minal Patel PhD [99], Verity Hill [90], Ms Claire M Bewshea MSc [91], Prof Sian Ellard FRCPath [73, 91], Dr Cressida Auckland FRCPath [73], Dr Ian Harrison [66], Dr Chloe Bishop [66], Dr Vicki Chalker [66], Dr Alex Richter PhD [85], Dr Andrew Beggs PhD [85], Dr Angus Best PhD [86], Dr Benita Percival PhD [86], Dr Jeremy Mirza PhD [86], Dr Oliver Megram PhD [86], Dr Megan Mayhew PhD [86], Dr Liam Crawford PhD [86], Dr Fiona Ashcroft PhD [86], Dr Emma Moles-Garcia PhD [86], Dr Nicola Cumley PhD [86], Mr Richard Hopes [64], Dr Patawee Asamaphan PhD [48], Mr Marc O Niebel MSc [48], Prof Rory N Gunson PhD FRCPath [100], Dr Amanda Bradley PhD [52], Dr Alasdair Maclean PhD [52], Dr Guy Mollett MBChB [52], Dr Rachel Blacow MBChB [52], Mr Paul Bird MSc [16], Mr Thomas Helmer [16], Miss Karlie Fallon [16], Dr Julian Tang [16], Dr Antony D Hale MBBS [49], Dr Louissa R Macfarlane-Smith PhD [49], Katherine L Harper MBiol [49], Miss Holli Carden MSc [49], Dr Nicholas W Machin MSc [45, 64], Ms Kathryn A Jackson MSc [92], Dr Shazaad S Y Ahmad MSc [45, 64], Dr Ryan P George PhD [45], Dr Lance Turtle PhD MRCP [92], Mrs Elaine O'Toole BSc [43], Mrs Joanne Watts BSc [43], Mrs Cassie Breen BSc [43], Mrs Angela Cowell MSc [43], Ms Adela Alcolea-Medina [32, 96], Ms Themoula Charalampous MSc [12, 42], Amita Patel [11], Dr Lisa J Levett PhD [35], Dr Judith Heaney PhD [35], Dr Aileen Rowan PhD [39], Prof Graham P Taylor DSc [39], Dr Divya Shah PhD [30], Miss Laura Atkinson MSc [30], Mr Jack CD Lee MSc [30], Mr Adam P Westhorpe BSc [82], Dr Riaz Jannoo PhD [82], Dr Helen L Lowe PhD [82], Miss Angeliki Karamani MSc [82], Miss Leah Ensell BSc [82], Mrs Wendy Chatterton MSc [35], Miss Monika Pusok MSc [35], Mrs Ashok Dadrah MSc [75], Miss Amanda Symmonds MSc [75], Dr Graciela Sluga MD/MSC [44], Dr Zoltan Molnar PhD [72], Mr Paul Baker MD [79], Prof Stephen Bonner [79], Ms Sarah Essex [79], Dr Edward Barton MD [56], Ms Debra Padgett BSc [56], Ms Garren Scott BSc [56], Ms Jane Greenaway MSc [57], Dr Brendan AI Payne MD [50], Dr Shirelle Burton-Fanning MD [50], Dr Sheila Waugh MD [50], Dr Veena Raviprakash MD [17], Ms Nicola Sheriff BSc [17], Ms Victoria Blakey BSc [17], ms Lesley-Anne Williams BSc [17], Dr Jonathan Moore MD [27], Ms Susanne Stonehouse BSc [27], Dr Louise Smith [55], Dr Rose K Davidson PhD [89], Dr Luke Bedford [26], Dr Lindsay Coupland PhD [54], Ms Victoria Wright BSc [18], Dr Joseph G Chappell PhD [97], Dr Theocharis Tsoleridis PhD [97], Prof Jonathan Ball PhD [97], Mrs Manjinder Khakh [15], Dr Vicki M Fleming PhD [15], Dr Michelle M Lister PhD [15], Dr Hannah C Howson-Wells PhD [15], Dr Louise Berry [15], Dr Tim Boswell [15], Dr Amelia Joseph [15], Dr Iona Willingham [15], Dr Nichola Duckworth [60], Dr Sarah Walsh [60], Dr Emma Wise PhD [2, 111], Dr Nathan Moore PhD [2, 111], Miss Matilde Mori BSc [2, 108, 111], Dr Nick Cortes MRCP FRCPath [2, 111], Dr Stephen Kidd PhD [2, 111], Dr Rebecca Williams BMBS [33], Laura Gifford MSc [69], Miss Kelly Bicknell [61], Dr Sarah Wyllie [61], Miss Allyson Lloyd [61], Mr Robert Impey MSc [61], Ms Cassandra S Malone MSc [6], Mr Benjamin J Cogger BSc [6], Nick Levene MSc [62], Lynn Monaghan [62], Dr Alexander J Keeley MRCP [93], Dr David G Partridge FRCP FRCPath [78, 93], Dr Mohammad Raza [78, 93], Dr Cariad Evans [78, 93] and Dr Kate Johnson [78, 93].

**Sequencing and analysis:**
Emma Betteridge BSc [99], Ben W Farr BSc [99], Scott Goodwin MSc [99], Dr Michael A Quail PhD [99], Carol Scott [99], Lesley Shirley MSc [99], Scott AJ Thurston BSc [99], Diana Rajan MSc [99], Dr Iraad F Bronner PhD [99], Louise Aigrain PhD [99], Dr Nicholas M Redshaw PhD [99], Dr Stefanie V Lensing PhD [99], Shane McCarthy [99], Alex Makunin [99], Dr Carlos E Balcazar PhD [90], Dr Michael D Gallagher PhD [90], Dr Kathleen A Williamson PhD [90], Thomas D Stanton BSc [90], Ms Michelle L Michelsen BSc [91], Ms Joanna Warwick-Dugdale BSc [91], Dr Robin Manley Ph.D. [91], Ms Audrey Farbos MSc [91], Dr James W Harrison Ph.D. [91], Dr Christine M Sambles Ph.D. [91], Dr David J Studholme PhD. [91], Dr Angie Lackenby [66], Dr Tamyo Mbisa [66], Dr Steven Platt [66], Mr Shahjahan Miah [66], Dr David Bibby [66], Dr Carmen Manso [66], Dr Jonathan Hubb [66], Dr Gavin Dabrera [66], Dr Mary Ramsay [66], Dr Daniel Bradshaw [66], Dr Ulf Schaefer [66], Dr Natalie Groves [66], Dr Eileen Gallagher [66], Dr David Lee [66], Dr David Williams [66], Dr Nicholas Ellaby [66], Hassan Hartman [66], Nikos Manesis [66], Vineet Patel [66], Juan Ledesma [67], Ms Katherine A Twohig [67], Dr Elias Allara [64, 88], Ms Clare Pearson [64, 88], Mr Jeffrey K. J. Cheng MSc [94], Dr Hannah E. Bridgewater PhD [94], Ms Lucy R. Frost BSc [94], Ms Grace Taylor-Joyce BSc [94], Dr Paul E Brown PhD [94], Dr Lily Tong PhD [48], Ms Alice Broos BSc [48], Mr Daniel Mair BSc [48], Mrs Jenna Nichols BSc [48], Dr Stephen N Carmichael PhD [48], Dr Katherine L Smollett PhD [40], Dr Kyriaki Nomikou PhD [48], Dr Elihu Aranday-Cortes PhD/DVM [48], Ms Natasha Johnson BSc [48], Dr Seema Nickbakhsh PhD [48, 68], Dr Edith E

Vamos PhD [92], Dr Margaret Hughes PhD [92], Dr Lucille Rainbow PhD [92], Mr Richard Eccles MSc [92], Ms Charlotte Nelson MSc [92], Dr Mark Whitehead PhD [92], Dr Richard Gregory PhD [92], Mr Matthew Gemmell MSc [92], Ms Claudia Wierzbicki BSc [92], Ms Hermione J Webster BSc [92], Ms Chloe L Fisher MSc [28], Mr Adrian W Signell BSc [20], Dr Gilberto Betancor PhD [20], Mr Harry D Wilson BSc [20], Dr Gaia Nebbia PhD FRCPath [12], Dr Flavia Flaviani PhD [31], Mr Alberto C Cerda MSc [96], Ms Tammy V Merrill MSc [96], Rebekah E Wilson MSc [96], Mr Marius Cotic MSc [82], Miss Nadua Bayzid BSc [82], Dr Thomas Thompson PhD [72], Dr Erwan Acheson PhD [72], Prof Steven Rushton PhD [51], Prof Sarah O'Brien PhD [51], David J Baker BEng [70], Steven Rudder [70], Alp Aydin MSci [70], Dr Fei Sang PhD [18], Dr Johnny Debebe PhD [18], Dr Sarah Francois PhD [23], Dr Tetyana I Vasylyeva DPhil [23], Dr Marina Escalera Zamudio PhD [23], Mr Bernardo Gutierrez MSc [23], Dr Angela Marchbank BSc [10], Joshua Maksimovic FD [9], Karla Spellman FD [9], Kathryn McCluggage Msc [9], Dr Mari Morgan PhD [69], Robert Beer BSc [9], Safiah Afifi BSc [9], Trudy Workman HNC [10], William Fuller BSc [10], Catherine Bresner Bsc [10], Dr Adrienn Angyal PhD [93], Dr Luke R Green PhD [93], Dr Paul J Parsons PhD [93], Miss Rachel M Tucker MSc [93], Dr Rebecca Brown PhD [93] and Mr Max Whiteley PhD [93]

**Software and analysis tools:**
James Bonfield BSc [99], Dr Christoph Puethe [99], Mr Andrew Whitwham BSc [99], Jennifier Liddle [99], Dr Will Rowe PhD [41], Dr Igor Siveroni PhD [39], Dr Thanh Le-Viet PhD [70] and Amy Gaskin MSc [69].

**Visualisation:**
Dr Rob Johnson PhD [39].

**1** Barking, Havering and Redbridge University Hospitals NHS Trust, **2** Basingstoke Hospital, **3** Belfast Health & Social Care Trust, **4** Betsi Cadwaladr University Health Board, **5** Big Data Institute, Nuffield Department of Medicine, University of Oxford, **6** Brighton and Sussex University Hospitals NHS Trust, **7** Cambridge Stem Cell Institute, University of Cambridge, **8** Cambridge University Hospitals NHS Foundation Trust, **9** Cardiff and Vale University Health Board, **10** Cardiff University, **11** Centre for Clinical Infection & Diagnostics Research, St. Thomas' Hospital and Kings College London, **12** Centre for Clinical Infection and Diagnostics Research, Department of Infectious Diseases, Guy's and St Thomas' NHS Foundation Trust, **13** Centre for Enzyme Innovation, University of Portsmouth (PORT), **14** Centre for Genomic Pathogen Surveillance, University of Oxford, **15** Clinical Microbiology Department, Queens Medical Centre, **16** Clinical Microbiology, University Hospitals of Leicester NHS Trust, **17** County Durham and Darlington NHS Foundation Trust, **18** Deep Seq, School of Life Sciences, Queens Medical Centre, University of Nottingham, **19** Department of Infection Biology, Faculty of Infectious & Tropical Diseases, London School of Hygiene & Tropical Medicine, **20** Department of Infectious Diseases, King's College London, **21** Department of Microbiology, Kettering General Hospital, **22** Departments of Infectious Diseases and Microbiology, Cambridge University Hospitals NHS Foundation Trust; Cambridge, UK, **23** Department of Zoology, University of Oxford, **24** Division of Virology, Department of Pathology, University of Cambridge, **25** East Kent Hospitals University NHS Foundation Trust, **26** East Suffolk and North Essex NHS Foundation Trust, **27** Gateshead Health NHS Foundation Trust, **28** Genomics Innovation Unit, Guy's and St. Thomas' NHS Foundation Trust, **29** Gloucestershire Hospitals NHS Foundation Trust, **30** Great Ormond Street Hospital for Children NHS Foundation Trust, **31** Guy's and St. Thomas' BRC, **32** Guy's and St. Thomas' Hospitals, **33** Hampshire Hospitals NHS Foundation Trust, **34** Health Data Research UK Cambridge, **35** Health Services Laboratories, **36** Heartlands Hospital, Birmingham, **37** Hub for Biotechnology in the Built Environment, Northumbria University, **38** Imperial College Hospitals NHS Trust, **39** Imperial College London, **40** Institute of Biodiversity, Animal Health & Comparative Medicine, **41** Institute of Microbiology and Infection, University of Birmingham, **42** King's College London, **43** Liverpool Clinical Laboratories, **44** Maidstone and Tunbridge Wells NHS Trust, **45** Manchester University NHS Foundation Trust, **46** Microbiology Department, Wye Valley NHS Trust, Hereford, **47** MRC Biostatistics Unit, University of Cambridge, **48** MRC-University of Glasgow Centre for Virus Research, **49** National Infection Service, PHE and Leeds Teaching Hospitals Trust, **50** Newcastle Hospitals NHS Foundation Trust, **51** Newcastle University, **52** NHS Greater Glasgow and Clyde, **53** NHS Lothian, **54** Norfolk and Norwich University Hospital, **55** Norfolk County Council, **56** North Cumbria Integrated Care NHS Foundation Trust, **57** North Tees and Hartlepool NHS Foundation Trust, **58** Northumbria University, **59** Oxford University Hospitals NHS Foundation Trust, **60** PathLinks, Northern Lincolnshire & Goole NHS Foundation Trust, **61** Portsmouth Hospitals University NHS Trust, **62** Princess Alexandra Hospital Microbiology Dept., **63** Public Health Agency, **64** Public Health England, **65** Public Health England, Clinical Microbiology and Public Health Laboratory, Cambridge, UK, **66** Public Health England, Colindale, **67** Public Health England, Colindale, **68** Public Health Scotland, **69** Public Health Wales NHS Trust, **70** Quadram Institute Bioscience, **71** Queen Elizabeth Hospital, **72** Queen's University Belfast, **73** Royal Devon and Exeter NHS Foundation Trust, **74** Royal Free NHS Trust, **75** Sandwell and West Birmingham NHS Trust, **76** School of Biological Sciences, University of Portsmouth (PORT), **77** School of Pharmacy and Biomedical Sciences, University of Portsmouth (PORT), **78** Sheffield Teaching Hospitals, **79** South Tees Hospitals NHS Foundation Trust, **80** Swansea University, **81** University Hospitals Southampton NHS Foundation Trust, **82** University College London, **83** University Hospital Southampton NHS Foundation Trust, **84** University Hospitals Coventry and Warwickshire, **85** University of Birmingham, **86** University of Birmingham Turnkey Laboratory, **87** University of

Brighton, **88** University of Cambridge, **89** University of East Anglia, **90** University of Edinburgh, **91** University of Exeter, **92** University of Liverpool, **93** University of Sheffield, **94** University of Warwick, **95** University of Cambridge, **96** Viapath, Guy's and St Thomas' NHS Foundation Trust, and King's College Hospital NHS Foundation Trust, **97** Virology, School of Life Sciences, Queens Medical Centre, University of Nottingham, **98** Wellcome Centre for Human Genetics, Nuffield Department of Medicine, University of Oxford, **99** Wellcome Sanger Institute, **100** West of Scotland Specialist Virology Centre, NHS Greater Glasgow and Clyde, **101** Department of Medicine, University of Cambridge, **102** Ministry of Health, Sri Lanka, **103** NIHR Health Protection Research Unit in HCAI and AMR, Imperial College London, **104** North West London Pathology, **105** NU-OMICS, Northumbria University, **106** University of Kent, **107** University of Oxford, **108** University of Southampton, **109** University of Southampton School of Health Sciences, **110** University of Southampton School of Medicine, **111** University of Surrey, **112** Warwick Medical School and Institute of Precision Diagnostics, Pathology, UHCW NHS Trust, **113** Wellcome Africa Health Research Institute Durban and **114** Wellcome Genome Campus.
